# Salamander-like tail regeneration in the West African lungfish

**DOI:** 10.1101/2020.02.12.946319

**Authors:** Kellen Matos Verissimo, Louise Neiva Perez, Aline Cutrim Dragalzew, Gayani Senevirathne, Sylvain Darnet, Wainna Renata Barroso Mendes, Ciro Ariel dos Santos Neves, Erika Monteiro dos Santos, Cassia Nazare de Sousa Moraes, Ahmed Elewa, Neil Shubin, Nadia Belinda Froebisch, Josane de Freitas Sousa, Igor Schneider

## Abstract

Salamanders, frog tadpoles, and diverse lizards have the remarkable ability to regenerate tails. Paleontological data suggests that this capacity is plesiomorphic, yet when the developmental and genetic architecture of tail regeneration arose is poorly understood. Here we show morphological and molecular hallmarks of tetrapod tail regeneration in the West African lungfish *Protopterus annectens*, a living representative of the sister group of tetrapods. As in salamanders, lungfish tail regeneration occurs *via* formation of a proliferative blastema and restores original structures, including muscle, skeleton and spinal cord. In contrast to lizards and similar to salamanders and frogs, lungfish regenerate spinal cord neurons and reconstitute dorsoventral patterning of the tail. Similar to salamander and frog tadpoles, *Shh* is required for lungfish tail regeneration. Through RNA-seq analysis of uninjured and regenerating tail blastema, we show that the genetic program deployed during lungfish tail regeneration maintains extensive overlap with that of tetrapods, with the upregulation of genes and signaling pathways previously implicated in amphibian and lizard tail regeneration. Furthermore, the lungfish tail blastema showed marked upregulation of genes encoding post-transcriptional RNA processing components and transposon-derived genes. Our results show that developmental processes and genetic program of tetrapod tail regeneration were present at least near the base of the sarcopterygian clade and establish the lungfish as a valuable research system for regenerative biology.

## 1. Introduction

In living tetrapods, the capacity to regenerate tails is present in salamanders, frog tadpoles, and some lizards. Paleontological evidence has shown salamander-like tail regeneration in lepospondyl microsaurs, tetrapods from the early Carboniferous and early Permian [1]. Whether salamander-like tail regeneration capacity was the early condition of this group remains uncertain. Given their phylogenetic position as the extant sister group to tetrapods, lungfish might hold the key to our understanding of the evolution of regenerative capacities in tetrapods.

Lungfishes are capable of regenerating complete tails as adults. This remarkable capacity was first reported nearly 150 years ago [2] and documented in the laboratory half a century ago [3]. Since then, despite its potential as a model research system for regenerative medicine and important phylogenetic position, no reports on lungfish tail regeneration have followed, likely due to the logistical difficulties in obtaining and housing lungfish as laboratory animals.

In contrast, decades of studies have and continue to shed light on the morphological and molecular processes of tail regeneration in tetrapods. In salamanders and frog tadpoles, soon after tail amputation, a wound epithelium forms and covers the injured site [4]. The thickening of the wound epidermis gives rise to the apical epithelial cap (AEC) and, at this point, patterns of tail regeneration diverge between salamanders and frog tadpoles. In salamanders, undifferentiated cells accumulate at the amputation site, giving rise to the blastema [4]. In tadpoles, a mass of undifferentiated cells accumulates around the neural ampulla and notochord tip, forming a blastema-like cell population, termed regeneration bud [5]. In salamanders, dedifferentiated cells play a significant role in restoring muscle tissue [6], whereas in tadpoles, this is achieved chiefly *via* proliferation of Pax7+ satellite cells [7,8]. Ultimately, frog tadpoles and salamanders regenerate tails with high fidelity, redeploying all tissue types and anatomical patterns of their original tail.

Lizards are the only amniotes capable of tail regeneration through wound healing and blastema formation. However, instead of regenerating the spinal cord and associated skeleton, a simple ependymal tube composed of neuroglia develops enclosed by an unsegmented cartilage tube [9,10]. The new ependymal tube and tail skeleton in lizards fail to recapitulate the embryonic dorsoventral expression pattern of genes responsible for establishing roof plate, floor plate, and the lateral domains of the neural tube. Specifically, floor plate markers *Shh* and *FoxA2* in lizards are expressed along the entire regenerating ependymal tube, which consequently acquires a floor plate identity [11].

Despite the differences in the regenerative process in amphibians and amniotes, molecular studies have revealed broad similarities in the genetic program of tail regeneration. Studies in frog tadpoles have identified a sequence of major molecular events leading to successful tail regeneration. In summary, upon injury, TGF-β signaling is required for the formation of the wound epidermis [12]. Extracellular matrix (ECM) remodeling, reactive oxygen species (ROS) signaling and inflammation are also among the first responses to tail injury [13]. Regeneration-organizing cells relocate to the wound epithelium and express Wnts and Fgfs [14]. As the regeneration bud forms and tail outgrowth occurs, additional signaling pathways are deployed, such as Bmp, Egf, Shh, Tgf-β and Notch pathways, among others [15, 16, 7]. Proliferating blastema cells show high expression levels of *Il11, Cse1l* and *L1td1-like* [17] and require histone deacetylases [18] and Hyaluronan, an ECM component [19] and *Il11* [20]. Likewise, during salamander tail regeneration, Wnt, Fgf, and Tgf-β pathways [16,21], ion channels [22], Shh signaling [21,23], Egf, Notch, and other signaling pathways [16] are required. Regenerating spinal cords also display upregulation of genes linked to immune and inflammatory response, ECM remodeling, and genes encoding morphogens such as Shh, Bmps, Wnts, and Fgfs [24]. Finally, morpholino-mediated knockdown of Marcks-like protein (Mlp), an extracellular protein, blocks tail regeneration [25].

The molecular profile of lizard tail regeneration has also been examined in recent years. In the common wall lizard (*Podarcis muralis*), genes exclusively upregulated in the regenerating tail include those coding for growth factors (Wnts, Shh, Bmps, Fgfs) and those within the broad categories of ECM, inflammation and immunity, metabolism and cytoskeleton [26]. Similarly, tail regeneration in geckos shows enrichment for peptides involved in immune response, ECM remodeling, Fgfs and Bmps [27] and involves ROS signaling [28], Tgf-β signaling [29], and Fgf signaling [30]. In the green anole lizard (*Anolis carolinensis*), the regenerating tail is enriched for the gene ontology (GO) categories of wound response, immune response, hormonal regulation and embryonic morphogenesis [31].

In sum, current molecular data has begun to reveal the broad genetic architecture of tetrapod tail regeneration, yet the evolutionary origins of this regenerative program remain unclear. Leveraging the phylogenetic position of lungfish as an outgroup of tetrapods we sought to characterize the morphological and genetic profile of lungfish tail regeneration. We found lungfish tail regeneration proceeded in a salamander-like manner, with the formation of a proliferative blastemal cell population and the restoration of original tail structures including muscle, spinal cord, and tail fin skeleton. Lungfish tail skeletal and spinal cord tissues regenerated neurons and proper dorsoventral patterning. Shh signaling, fundamental for amphibian tail dorsoventral patterning and regeneration, is also required for lungfish tail regeneration. RNA-seq analysis revealed that lungfish deploy a blastemal genetic program similar to that reported in tetrapods. Interestingly, lungfish tail blastema showed marked upregulation of transposon-derived genes and components of post-transcriptional RNA processing. Our findings suggest that salamander-like tail regeneration was present in the sarcopterygian ancestor of tetrapods and lungfish.

## 2. Methods

### (a) Animals and surgical procedures

Thirty juvenile West African Lungfish (*Protopterus annectens*) ranging from 15 to 35 cm in length, were acquired through the pet trade and housed at the Universidade Federal do Para. Specimens were kept in individual 4 L gallon tanks in dechlorinated tap water at 24-28 °C with aeration and biological and mechanical filtration. Lungfish were anesthetized in 0.1% MS-222 (Sigma Aldrich). The position of amputation was determined as follows: the distance from the snout to the attachment point of the pelvic fin was measured in centimeters and divided by 1.75. The resulting number was then used as the distance from the attachment point of the pelvic fin to the point of tail amputation. Samples were labeled as ‘uninjured’ or ‘blastema’ and embedded in Tissue Tek O.C.T compound (Sakura Finetek), or stored in RNAlater (Sigma-Aldrich) at −80 °C temperature for RNA extraction, frozen on dry ice for histology and *in situ* hybridization (stored at −80 °C), or fixed in 4% paraformaldehyde (PFA) overnight at 4 °C for section-immunohistochemistry.

### (b) External morphology and histology of tail fin regeneration

Tails from animals under anesthesia were photographed at 1 day post-amputation (dpa) and weekly to document the changes on external morphology during regeneration. For histology, frozen tissues were allowed to adjust to cryostat temperature (−20 °C) for 30 min. Next, 20 μm longitudinal sections were obtained and placed on ColorFrost Plus microscope slides (Thermo Fisher Scientific). Sections were fixed in 3% PFA for 5 min, rinsed twice in 0.1M PBS, and dehydrated in graded ethanol series (70, 95 and 100%) for 2 min each. Slides were stored at −80 °C. Sections were stained with hematoxylin (Sigma-Aldrich) and eosin (Sigma-Aldrich) and imaged on an SMZ1000 stereoscope (Nikon). Tails were cleared and stained as described previously [32]. In total, histological sections were obtained from 6 animals, and tails from 5 animals were used for clearing and staining.

### (c) Inhibition of Shh signaling

Cyclopamine (Selleckchem, cat. number S1146) dissolved in DMSO was added to the aquarium water at a final concentration of 1 μg/ml. Control group was treated with DMSO at a final concentration of 0.1%. In both DMSO-only (n = 3) and cyclopamine-treated (n = 3) groups, aquarium water was changed daily with fresh cyclopamine or DMSO-only solution for 6 weeks. Animals were photographed and measurements of total tail length were taken weekly.

### (d) Cell proliferation assay and immunohistochemistry

5-bromo-2-deoxyuridine (BrdU) was injected intraperitoneally (80 mg per kg of body weight) into anesthetized lungfish 24 h before tail tissue collection to observe cell proliferation. Overnight-fixed tissues were transferred to 30% sucrose, flash-frozen in OCT blocks; longitudinal and transverse sections (20 μM thickness) were obtained as described in the preceding section. Sections were permeabilized in 2N HCl solution at 37 °C for 15 minutes, followed by washes in 0.1M borate buffer and in PBS tween (0.1% tween in 0.01M PBS). Treatment with 0.1% trypsin at 37 °C for 15 minutes was performed and followed by a wash in PBS tween. Unspecific labeling was blocked with 5% normal goat serum diluted in 0.01M PBS with 0.5% Tween and 1% bovine serum albumin for 1 h at room temperature. Next, sections were incubated with mouse anti-BrdU primary antibody (1:200, Sigma-Aldrich, cat. number B8434) in 0.01 M PBS with 1% bovine serum albumin and 0.5% Tween overnight at 4 °C. On the following day, sections were incubated with the Alexa 488 conjugated goat anti-mouse secondary antibody (1:400, ThermoFisher Scientific, cat. number A-11001) for 2 h at room temperature and slides were mounted and counterstained with Fluoromount with DAPI (ThermoFisher Scientific). For immunohistochemistry, βIII-Tubulin immunostaining (mouse monoclonal, 1:500, Sigma-Aldrich, cat. number ab78078) was performed using the same procedure except for HCL and trypsin treatments, followed by incubation with the Alexa Fluor 594 goat anti-mouse secondary antibody (1: 1000, ThermoFisher Scientific, cat. number A-11005).

### (e) Library preparation and Illumina sequencing

For transcriptome sequencing, total RNA from tail tissues was extracted using TRIzol Reagent (Thermo Fisher). A two-step protocol, with the RNeasy Mini Kit (Qiagen) and DNase I treatment (Qiagen), were used to purify the RNAs and remove residual DNA. mRNA sequencing libraries were constructed using the NEXTflex^®^ Rapid Directional qRNA-Seq™ Kit (Illumina). Lungfish reference transcriptomes and transcript abundance estimation were obtained from the sequencing of three biological replicates of blastemas at 14 dpa and three biological replicates of uninjured tail tissue, performed on an Illumina 2500 HiSeq platform with 150 bp paired-end reads. Reads from 6 additional runs of other regenerating tail stages were used only to help produce a comprehensive *de novo* lungfish reference transcriptome assembly.

### (f) Bioinformatic analysis

The West African lungfish reference transcriptome was assembled *de novo* using Trinity-v2.9.0 with default parameters [33]. The transcriptome assembly was subjected to the EvidentialGene pipeline for greater accuracy of gene set prediction [34]. For each run, all read datasets were mapped to reference transcriptomes using CLC genomic workbench with default parameters (Qiagen). Expression data, measured by transcript count, were summed by human homolog gene cluster (HHGC) using a custom bash script. As previously described [35], the HHGCs were defined by grouping transcripts with an e-value of 10^−3^ when compared by BLASTx against Human NCBI RefSeq database. For each HHGC, expression was calculated in transcripts per million (TPM) and mean TPM value between uninjured tails and tail blastemas were compared with a two-tailed t-test using the CLC genomic workbench with default parameters (Qiagen). A transcript or HHGC was deemed differentially expressed when its fold-change is greater than 2 or less than −2 and the FDR adjusted *P*-value < 0.05. A similarity matrix between samples wa*s* calculated using square root transformed TPM values for each HHGC, using Spearman rank correlation in Morpheus software (https://software.broadinstitute.org/morpheus). A list of enriched GO terms and overrepresented Reactome pathways was produced using WebGestalt 2019 [36]. Differentially expressed genes with false discovery rate (FDR) adjusted *P*-values smaller than 0.05 were ranked from highest to lowest fold change values, and the corresponding ranked list of gene symbols was used for the GO enrichment analysis. GO enriched categories or over-represented Reactome pathways were significant when *P* values were 0.05 or less. Venn diagrams were generated using BioVenn [37]. GO enrichment and pathway over-representation analyses were performed using WebGestalt 2019 web-based tool. Protein domains were identified using HMMER v3.2.1 (http://hmmer.org/) against the proteome generated using TransDecoder v5.3.0 (http://transdecoder.github.io).

### (g) *In situ* hybridization

Frozen sections (20 μm) from regenerating tails at 14 dpa (n = 3) were obtained on a Leica CM1850 UV cryostat and positioned on the Color Frost Plus microscope slides (Thermo Fisher). Sections were fixed as previously described [35] and stored at −80°C for hematoxylin and eosin staining or *in situ* hybridization. Riboprobe templates for *in situ* hybridization were produced by a two-round PCR strategy: first-round PCR produced specific fragments (400-500 bp) of selected genes, and in a second PCR a T7 promoter sequence was included at either 5’or 3’end of the fragments for the generation of templates for sense or anti-sense probes. The primers used were: *Col12a1* forward: 5’-<u>GGCCGCGG</u>TTGATGCTCCCATTTGGTTAG-3’ and reverse: 5’-<u>CCCGGGGC</u>GAAACCCAGGAACAAGAGGTC-3’, *Hmcn2* forward 5’-<u>GGCCGCGG</u>TTGAGCAGAACCAGCTTCATT-3’ and reverse 5’-<u>CCCGGGGC</u>TTAGTGGGGCAGACAATCAAC-3’, *Inhbb* forward 5’-<u>GGCCGCGG</u>CCGTGCTTGAACCACTAAAAA and reverse 5’-<u>CCCGGGGC</u>TTTGCAGAGACAGATGACGTG-3’, 3’-T7 universal 5’-AGGGATCCTAATACGACTCACTATAGGGCCCGGGGC-3’, 5’-T7 universal 5’-GAGAATTCTAATACGACTCACTATAGGGCCGCGG-3’ (linker sequences for annealing of the 3’ or 5’ T7 universal primer are underlined). The riboprobes were synthesized using T7 RNA polymerase (Roche) and DIG-labelling mix (Roche). *In situ*, hybridization was performed as previously described [35], using 375 ng of DIG-labelled riboprobe per slide. Slides were photographed on the Nikon Eclipse 80i microscope and the images were processed on the NIS-Element D4.10.1 program.

## 3. Results

### (a) Establishment of a proliferative blastemal cell population during lungfish tail regeneration

We evaluated regeneration in juvenile West African lungfish upon tail amputation (figure 1a). At 7 dpa, a wound epithelium covers the amputation site, and tail outgrowth is negligible. At 14 dpa, tail outgrowth is visible at the level of the midline. At 21 dpa, the regenerating tail reaches approximately 1 cm in length and the skin is highly pigmented. In the following weeks, the tail continues to extend and by 56 dpa it has nearly reached its length before amputation. Histological sections showed that at 1 dpa, a 1 to 2 cell layer wound epithelium covers the amputation site (figure 1b). At 7 dpa, the wound epithelium thickens, and a mass of mesenchymal cells accumulate subjacent to it, posterior to the severed postcaudal cartilage. At 21 dpa, the regenerated post caudal cartilage bar and ependymal tube are visible. At our latest experimental endpoint (60 dpa), the postcaudal cartilage was undergoing segmentation and cartilaginous neural and haemal arches and spines were visible (figure 1c).

**Figure 1.**
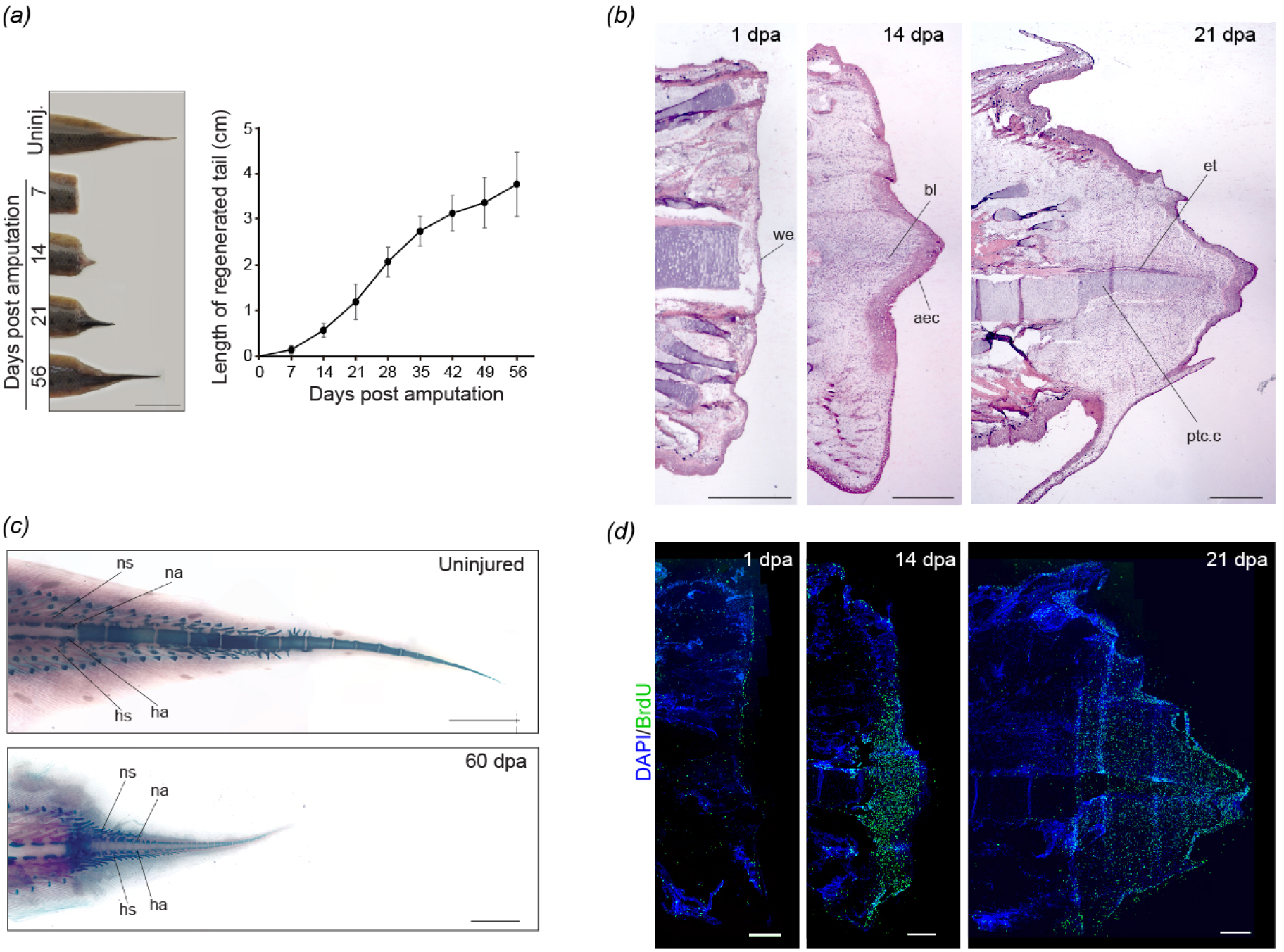
Morphological characterization of tail regeneration in the West African lungfish. (a) shows the progression of lungfish tail regeneration and the extent of growth up to 56 dpa. Vertical bars in the graph represent standard deviation. (b) histological sections of regenerating lungfish tail. (c) regeneration of skeletal elements of the tail at 60 dpa. (d) BrdU staining of proliferative cells during tail regeneration. we, wound epithelium; aec, apical epithelial cap; bl, blastema; et, ependymal tube; ptc.c, postcaudal cartilage; ns, neural spine; na, neural arch; hs, haemal spine; ha, haemal arch. Scale bars of 1 cm (a), 1 mm (b,d), 0.5 cm (c).

Next, we assessed cell proliferation during the first 3 weeks of tail regeneration. BrdU staining revealed that at 1 dpa, proliferating cells are mostly found in the wound epithelium. At 14 dpa, proliferating cells are found distal to the amputation plane in the region of the presumptive tail blastema. At 21 dpa, cell proliferation is observed posterior to the amputation site across the entire regenerated tail (figure 1d). Our results indicate that lungfish tail regeneration proceeds *via* morphological events similar to those involved in salamander tail regeneration, with the establishment of a wound epithelium, which thickens to form an AEC, the formation of a mass of proliferating blastemal cells, and restoration of original tail tissue organization.

### (b) Lungfish regeneration restores spinal cord neurons and original dorsoventral patterning of the original tail

To determine whether lungfish regenerating tails reestablish dorsoventral tail patterning and spinal cord neurogenesis, we examined transversal sections of lungfish uninjured and regenerating tails. We found that at 21 dpa, the regenerating ependymal tube is dorsally positioned relative to the notochord (figure 2a). A newly formed blood vessel ventral to the notochord and regenerating muscle are also visible. In addition, immunostaining for βIII-Tubulin revealed new spinal cord neurons forming as early as 28 dpa (figure 2b).

**Figure 2.**
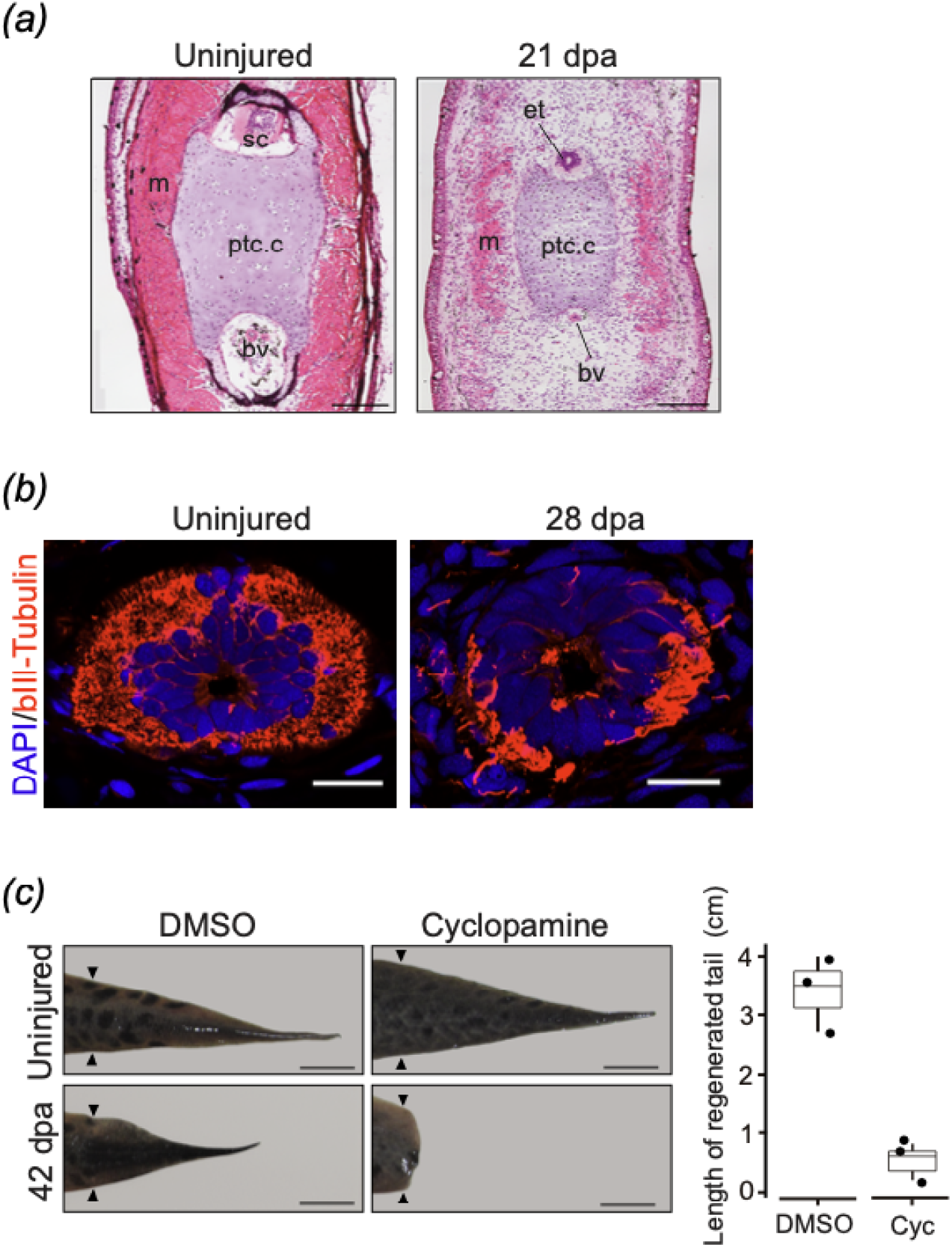
Establishment of dorsoventral organization and the requirement for Shh signaling during lungfish tail regeneration. (a) histological transversal sections of uninjured and 21 dpa regenerating tail. (b) immunostaining of DAPI and βIII-tubulin in uninjured and 28 dpa regenerating spinal cord (c) Effect of DMSO and cyclopamine treatment in tail regeneration. m, muscle; ptc.c, postcaudal cartilage; et, ependymal tube; bv, blood vessel. Scale bars of 1 mm (a - panoramic views - and b), 0.5 mm (a, enlarged view). Bars in graph represent standard deviation (c).

Shh is a key signaling molecule expressed in the floor plate of the ependymal tube in salamanders and the regenerating notochord in frog tadpoles, which is necessary for tail regeneration in both species [23,38]. To test its requirement for lungfish tail regeneration, we performed pharmacological inhibition of Shh signaling *via* administration of the cyclopamine. We found that in contrast to DMSO treatment (control group), continuous exposure to cyclopamine completely blocked lungfish tail regeneration, assessed at 42 dpa (figure 2c). Our results suggest that reestablishment of dorsoventral patterning, spinal cord neurogenesis and a requirement of Shh signaling might represent plesiomorphic features of tail regeneration.

### (c) Differential gene expression analysis of tail blastema versus uninjured tail

To identify genes differentially expressed in the tail blastema relative to uninjured tail tissue, we produced RNA-seq libraries from uninjured tail tissues and regenerating tails at 14 dpa, a stage when proliferative blastemal cells were identified. Principal component analysis showed two distinct clusters representing uninjured and blastemal tail samples, and Spearman correlation coefficients among biological replicas were greater than 0.78, corroborating the reproducibility of RNA-seq runs (electronic supplementary material, figure S1). DGE analysis of the lungfish uninjured *versus* 14 dpa tail revealed 1072 upregulated genes (FC > 2, FDR < 0.05). Among the upregulated gene dataset, we identified components of the various pathways previously associated to tail regeneration in tetrapods, including the Wnt pathways (*Wnt5a, Wnt5b, Axin2, Gsk3b, Ctnnb1*), Fgf (*Fgfr1*), Bmp (*Bmp1, Bmp4, Smad2*), Shh (*Ptch2, Gli2*), Notch (*Notch2*), Tgf-β (*Inhbb, Tgfbi*), Egf (*Vegfa, Megf6, Megf10*), ECM components and remodelers (*C1qtnf3, Col11a1, Fbn2, Mmp11, Adamts14*), Hyaluron pathway (*Hyal2, Has2*), immune and inflammatory response (*Il11*, *Mdk, Nfkbiz*) and stem cell maintenance (*Sox4*, *Sall4, Chd2*) (figure 3a and c). Some components of pathways previously involved in tetrapod tail regeneration were moderately upregulated (FC > 1.5, FDR < 0.05), including *Bmp2, Hif1a, Tgfb1, Tgfb2, Myc* and *Hdac7* (electronic supplementary material, table S1).

**Figure 3.**
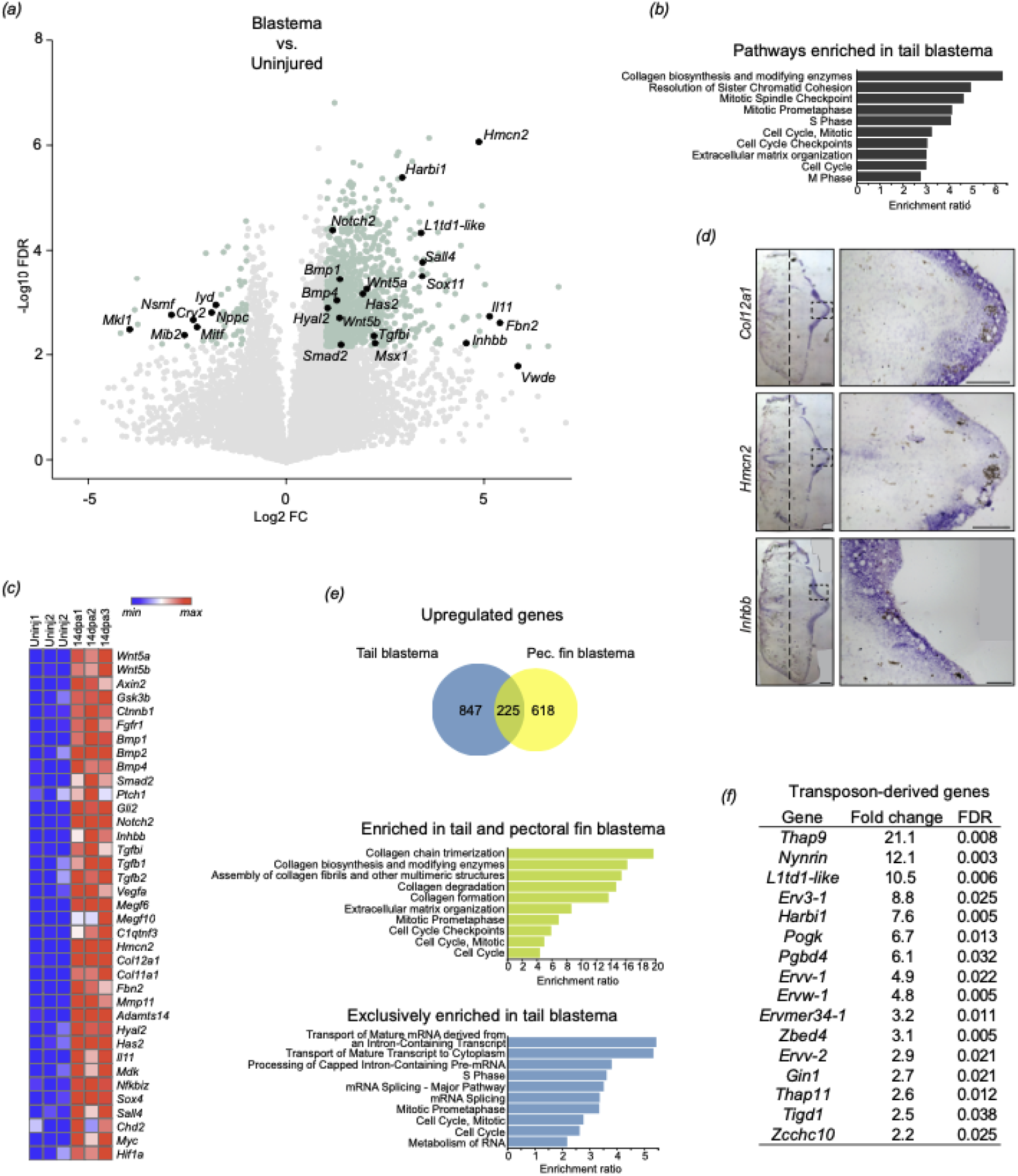
Upregulated genes and overrepresented pathways in lungfish tail blastema relative to uninjured tail. (a) Volcano plot showing differentially expressed genes in lungfish uninjured tail tissue and 14 dpa tail blastema (FDR < 0.05, FC > 2), Selected lungfish orthologs up or downregulated in the blastema are noted as black dots. (b) Pathways overrepresented in the tail blastema. (c) heatmap denoting subset of upregulated genes. (d) *in situ* hybridization of genes upregulated in the blastema (e) Area-proportional Venn diagram showing commonly upregulated genes in lungfish tail and pectoral fin datasets, enriched pathways in the shared tail and pectoral fin dataset, and pathways enriched exclusively on tail blastema. (f) Transposon-derived genes upregulated in the tail blastema. Scale bars of 1 mm (panoramic views) and 0.25 mm (enlarged view). In (c), “max” and “min” represent maximum and minimum expression levels of each gene.

GO enrichment analysis identified categories such as mitotic cell cycle phase transition, extracellular structure organization, and regulation of RNA metabolic processing (electronic supplementary material, figure S2). Likewise, pathway enrichment analysis revealed that the lungfish tail blastema shows a high overrepresentation of genes in collagen biosynthesis and modifying enzymes, pathways related to mitotic cell division, and ECM organization, consistent with the major events occurring at the blastema stage of tail regeneration (figure 3b). *In situ* hybridization in 14 dpa lungfish tails of 2 genes encoding ECM components (*Col12a1* and *Hmcn2*) and a gene encoding a member of the TGF-β family of cytokines (*Inhbb*) showed similar expression pattern, with anti-sense probe signal predominantly detected in the AEC (figure 3d), and no specific signal observed in sense-control probes (electronic supplementary material, figure S3).

*Il11*, a gene highly upregulated and required for tail regeneration in *Xenopus* tadpole, was also among the most highly upregulated in our dataset (FC = 34.95) (electronic supplementary material, table S1). The lungfish ortholog of *Mlp*, required for axolotl tail regeneration, showed moderate upregulation in lungfish tail and FDR value just above our cutoff (FC = 1.41, FDR = 0.06). The lungfish ortholog of *Vwde*, a gene highly expressed and required for axolotl limb regeneration and associated with successful frog tadpole tail regeneration [39], was highly upregulated in lungfish, however, with an FDR value above our cutoff (FC > 57.45, FDR = 0.08). Furthermore, *Angptl2* and *Egfl6*, identified recently as tail-specific AEC factors in frog tadpoles [40], are both upregulated in our lungfish tail blastema dataset (FC = 3.67 and 7.62, respectively) (electronic supplementary material, table S1). Interestingly, we also found 16 transposon-derived genes upregulated in the tail blastema (figure 3f), including *Ltd1-like*, a gene found enriched in the tail blastema of *Xenopus* tadpoles [17].

Finally, we examined a set of 10 genes recently reported to be expressed preferentially in the tail blastema of *frog* tadpoles relative to embryonic tail bud [17]. We found that 3 out of 10 genes were *Xenopus*-specific genes, and one gene was not contained in our annotated lungfish reference transcriptome. Of the 6 remaining, 3 were upregulated in our lungfish dataset, namely *Il11*, *Cse1l* (FC = 2.77), *L1td1* (FC = 10.52), and 1 gene, *cd200*, was upregulated with an FDR value above our 0.05 cutoff (FC = 5.94, FDR = 0.09). Taken together, our results identify general features of a genetic program of tail regeneration that may have been present in the last common ancestor of lungfish and tetrapods.

### (d) Genes enriched in lungfish tail blastema versus pectoral fin blastema

Next, we sought to compare our tail blastema dataset to previously published data on South American lungfish pectoral fin regeneration [32]. Comparison of 1072 upregulated genes in tail blastema to the 843 genes upregulated in pectoral fin blastema revealed an overlap of 225 genes. Reactome pathway enrichment analysis showed that this overlapping dataset included genes involved in collagen metabolism, ECM organization, and mitotic cell cycle, all of which represent categories commonly found in regenerating tissues (figure 3d). Interestingly, genes exclusively enriched in tail blastema relative to pectoral fin blastema were involved in pathways related to post-transcriptional RNA processing, including transport of mature transcript to the cytoplasm, processing of capped intron-containing pre-mRNA, mRNA splicing and metabolism of RNA (figure 3e). These results suggest that post-transcriptional RNA processing may play a more significant role in tail versus pectoral fin regeneration.

## 4. Discussion

Here we provided evidence of morphological and molecular hallmarks of tetrapod tail regeneration in the West African lungfish. In terms of morphology, lungfish tail regeneration was most similar to salamanders, featuring the formation of a highly proliferative wound epithelium, which thickened to form an AEC; a highly proliferative blastemal cell population; restoration of the proper dorsoventral pattern of the tail constituents; spinal cord neurogenesis; and requirement of Shh signaling. RNA-seq analysis of regenerating lungfish tail blastema revealed marked upregulation of signaling pathways previously linked to tail regeneration in tetrapods, such as Wnt, Fgf, Shh, Notch, Tgf-β and Egf, as well as ECM and inflammatory response. In addition to broad similarities, genes related to specific aspects of amphibian tail regeneration were also detected, such as the tail-specific AEC factors *Angptl2* and *Egfl6, Cse1l, L1td1-like*, and *Il11*.

Like salamanders, lungfish can regenerate paired appendages in addition to tails. When we compared enriched pathways in tail and paired-fin blastema, we found that pathways related to RNA processing were preferentially enriched in tails. It is possible that the greater complexity of cell types and stem and progenitor cell dynamics (especially in the nervous tissue) involved in tail *versus* paired-fin regeneration might account for an increased post-transcriptional RNA processing [41–43].

Interestingly, we also found evidence of upregulation of transposon-derived genes. Transposable elements may have played a role in the genome expansion in the lungfish [44], the Iberian ribbed newt *Pleurodeles waltl* [45], the axolotl [46] and the coelacanth [47–49]. In *P. waltl*, the transposable element family that expanded the most was the Harbinger transposon family, which has given rise to two vertebrate proteincoding genes, *Harbi1* [50] and *Naif1* [51]. In the coelacanth, Harbinger elements accounted for 4% of the genome and were shown to possess transcriptional and enhancer activities *in vivo* [48]. In our dataset, the lungfish *Harbi1* orthologue was among the highest differentially expressed transposon-derived transcripts. Future studies aimed at functionally evaluating the roles of transposon-derived genes such as *Harbi1* may uncover specific roles in development and regeneration.

Our results, together with recent paleontological findings, provide support for tail regeneration as a plesiomorphic trait present in the common ancestor of tetrapods and lungfish (figure 4). In this scenario, tail regeneration in lizards represents a derived character state, possibly with reemergence of regenerative capacity, where tail structures fail to recapitulate the tissue organization and composition seen in the uninjured tail. It is interesting to note that the fossil microsaur *Microbrachis* shows a pattern of tail regeneration that is very similar to what is seen in modern salamanders [1,52]. These fossils are members of the stem lineage of amniotes that lived in the Upper Carboniferous, about 300 million years ago [53]. Tail regeneration in these fossils has been previously discussed [54,55] and recently it was noted more concretely that the pattern of regeneration is indeed comparable to modern salamanders, in that vertebral elements are replaced, presumably with the associated spinal cord and musculature [1,52]. Moreover, vertebral centra are forming first in the regenerating tail of *Microbrachis*, before the associated neural arches, which is the reverse order of events seen in tail development and the same pattern seen in salamanders [52,56]. The morphological observations in fossil microsaurs suggest that at this point of evolutionary time, members of the stem lineage of amniotes were apparently still able to reactivate the positional information and tissue organization during tail regeneration. Given that the developmental pattern of lizard tail regeneration is significantly different from what is seen in amphibians and lungfish, and the phylogenetic distribution of non-regenerating species in other amniotes, we predict that tail regeneration re-evolved in lizards.

**Figure 4.**
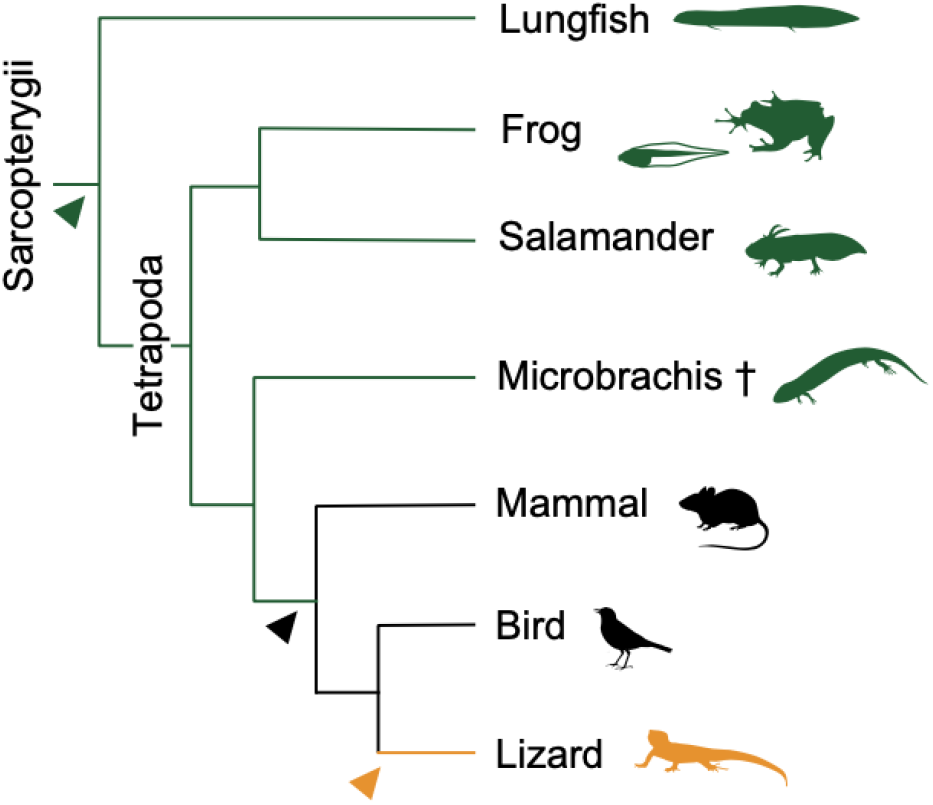
Hypothesis for the evolution of tail regeneration in sarcopterygians. Regeneration-incompetent lineages are shown in black, lineages with one or more regeneration-competent species are shown green, orange denotes *de novo* appearance of tail regeneration in Lepidosauria; green arrowhead indicates earliest occurrence of tail regeneration, black arrowhead indicates earliest loss, and orange arrowhead, reemergence. Cross signifies extinct taxon.

Our findings indicate that salamander-like tail regeneration was present in the common ancestor of lungfish and tetrapods, yet the origins of this ability may reside much earlier in evolutionary time, since amphioxus tail regeneration shares parallels with that seen in vertebrates [57]. Expanding the array of research species and searching for evidence of tail regeneration in fossil fish may help us better understand the evolutionary history of tail regeneration. Finally, our work underscores the importance of lungfish for our understanding of how tetrapod traits evolved and helps establish lungfish as an emerging model system that can inform both the history and mechanisms of regeneration.

## Ethics

All experimental procedures and animal care were conducted following the Ethics Committee for Animal Research at the Universidade Federal do Pará, under the approved protocol number 037-2015.

## Authors’ contributions

KMV, JFS, NHS, NBF and IS designed the research; KMV, JFS, ACD, WRBM, CASN, EMS, CNSM and IS performed regeneration assays; KMV, WBM, CNSM, GS and EMS, performed cell proliferation assay, immunohistochemistry, and pharmacological experiments; LNP, KMV, JFS, ACD, SD, NBF, AE and IS analyzed transcriptome data; IS supervised this work and wrote the manuscript with input from all authors. All authors gave final approval for publication and agreed to be held accountable for the work performed therein.

## Data accessibility

Sequence data that support the findings of this study have been deposited in GenBank with the following BioProject accession numbers: PRJNA491932, with accession number for uninjured (SRR7880018, SRR7880019 e SRR7880016), 14 dpa blastema (SRR7880020, SRR7880021 e SRR7880024) and additional libraries of tail blastemas at 1 dpa (SRR7880017, SRR7880022 e SRR7880023) and 21 dpa (SRR7880025, SRR7880026 e SRR7880027). The authors declare that all other relevant data supporting the findings of this study are available on request.

## Acknowledgments

We would like to thank Jamily Lima for help with illustrations. We also thank Thomas Stewart, Justin Lemberg and Patricia Schneider for insightful comments on the manuscript.

---

**Figure S1.**
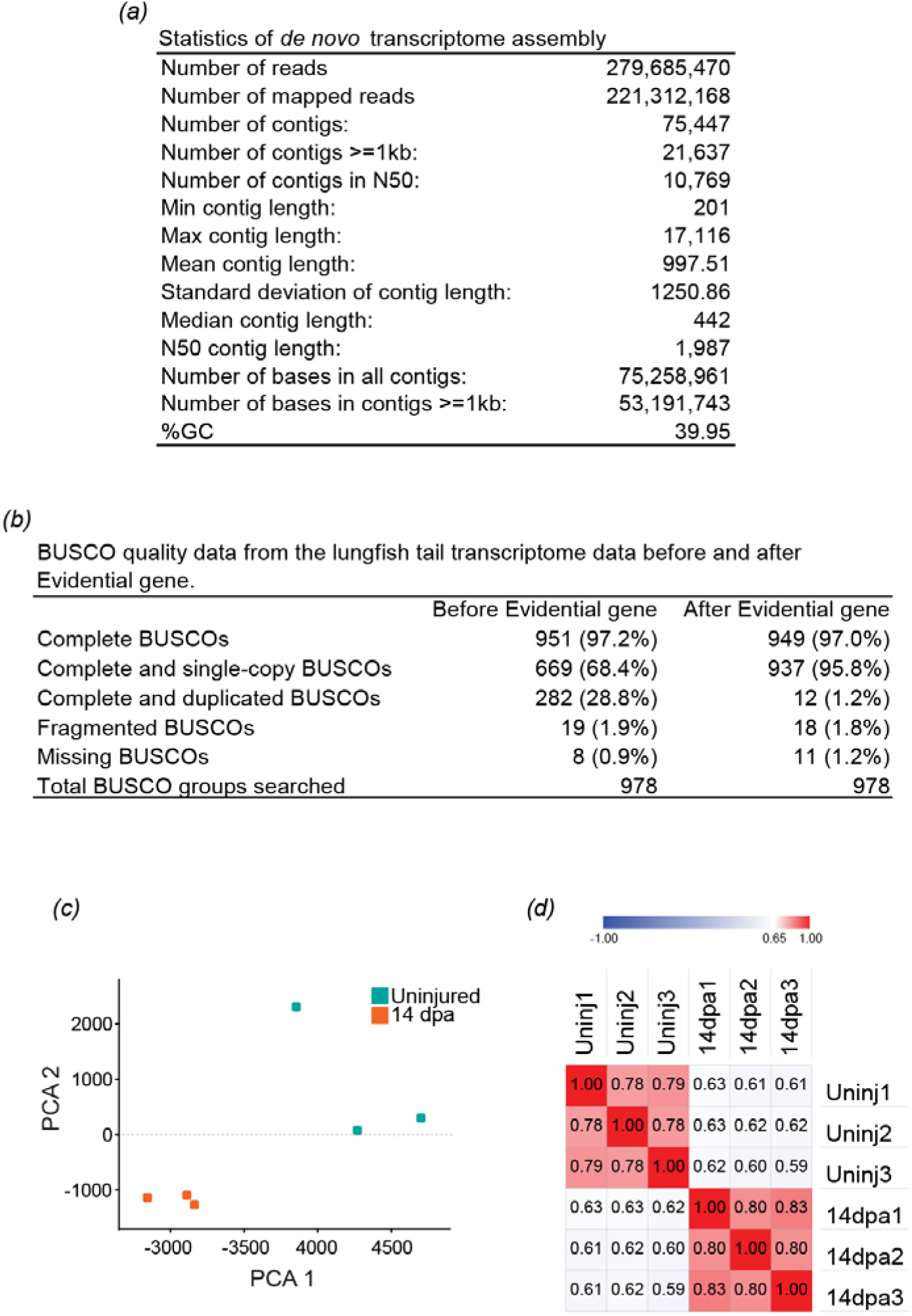
Statistics of the lungfish reference transcriptome and similarity among RNA-seq libraries. (a) Statistics or the lungfish reference transcriptome. (b) BUSCO assessment of transcriptome completeness. (c) Principal component analysis (PCA) of uninjured and 14 dpa lungfish tail libraries. (d) Spearman correlation matrix of uninjured and 14 dpa lungfish tail libraries.

**Figure S2.**
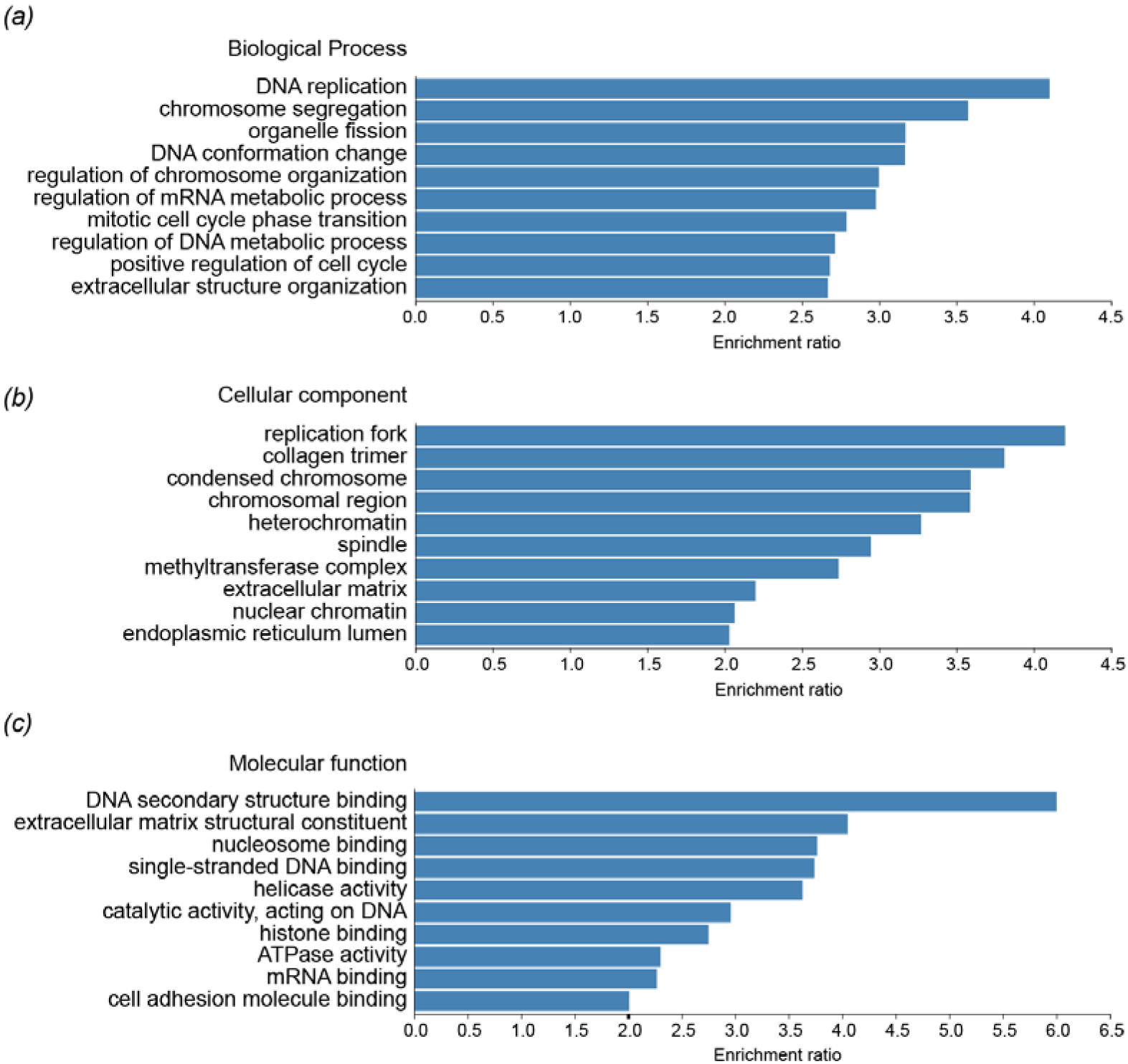
Gene ontology enrichment analysis of genes upregulated in the lungfish 14 dpa tail blastema as compared to uninjured tail. (a) Biological process. (b) Cellular component. (c) Molecular function.

**Figure S3.**
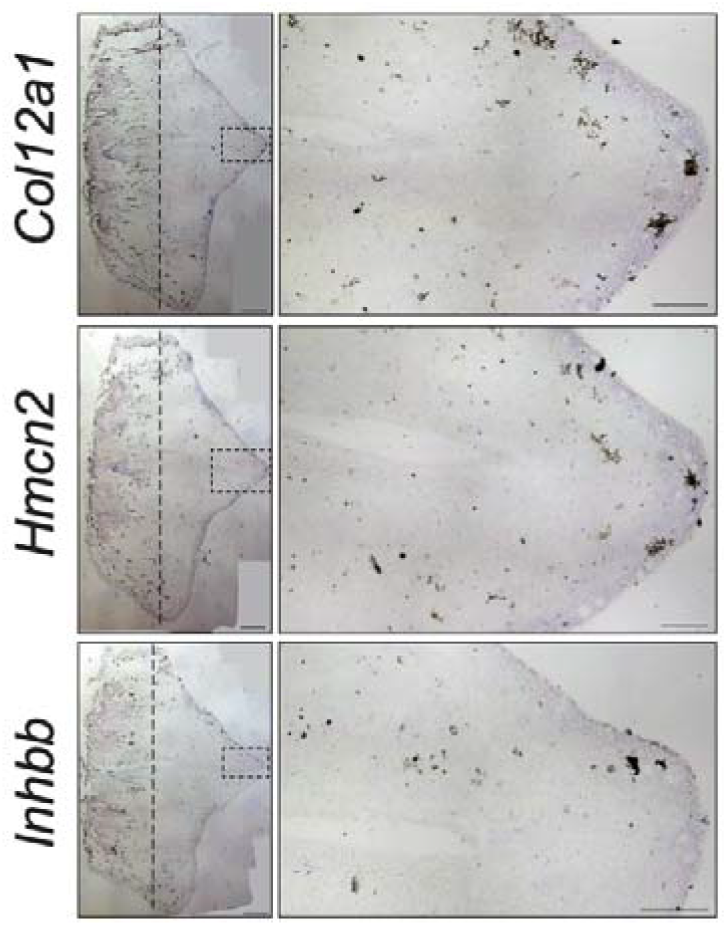
*In situ* hybridization of sense-control probes for select genes upregulated in the 14 dpa lungfish tail blastema relative to uninjured tail tissue. Scale bars of 1 mm (panoramic views) and 0.25 mm (enlarged view).

